# Dynamic compensation and homeostasis: a feedback control perspective

**DOI:** 10.1101/251298

**Authors:** Michel Fliess, Cédric Join

## Abstract

“Dynamic compensation” is a robustness property where a perturbed biological circuit maintains a suitable output [Karin O., Swisa A., Glaser B., Dor Y., Alon U. (2016). Mol. Syst. Biol., 12: 886]. In spite of several attempts, no fully convincing analysis seems now to be on hand. This communication suggests an explanation via “model-free control” and the corresponding “intelligent” controllers [Fliess M., Join C. (2013). Int. J. Contr., 86, 2228-2252], which are already successfully applied in many concrete situations. As a byproduct this setting provides also a slightly different presentation of homeostasis, or “exact adaptation,” where the working conditions are assumed to be “mild.” Several convincing, but academic, computer simulations are provided and discussed.

## 1. Introduction

In *systems biology, i.e*., an approach of growing importance to theoretical biology (see, *e.g*., Alon (2006); Klipp *et al*. (2016); Kremling (2012)), basic control notions, like feedback loops, are becoming more and more popular (see, *e.g*., Åström *et al*. (2008); Cowan *et al*. (2014); Cosentino *et al*. (2011); Del Vecchio *et al*. (2015)). This communication intends to show that a peculiar feedback loop permits to clarify the concept of *dynamic compensation* (*DC*) of biological circuits, which was recently introduced by Karin *et al*. (2016). DC is a robustness property. It implies, roughly speaking, that biological systems are able of maintaining a suitable output despite environmental fluctuations. As noticed by Karin *et al*. (2016) such a property arises naturally in physiological systems. The DC of blood glucose, for instance, with respect to variation in insulin sensitivity and insulin secretion is obtained by controlling the functional mass of pancreatic beta cells.

The already existing and more restricted *homeostasis*, or *exact adaptation*, deals only with constant reference trajectories, *i.e*., setpoints. It is achievable via an *integral* feedback (see, *e.g*., Alon *et al*. (1999); Briat *et al*. (2016); Miao *et al*. (2011); Stelling *et al*. (2004); Yi *et al*. (2000))

#### Remark 1.1.

*PIDs (see, e.g., Åström et al. (2008); O’Dwyer (2009)) read:*

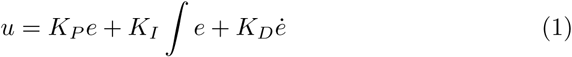

*where*

- *u, y, y* are respectively the control and output variables, and the reference trajectory*.
- *e* = *y − y* is the tracking error*,
- *K*_*P*_, *K*_*I*_, *K*_*D*_ ∈ ℝ *are the tuning gains*.

*To the best of our knowledge, they are, strangely enough, more or less missing in the literature on theoretical biology*,^1^ *although they lead to the most widely used control strategies in industry*.

*From K*_*P*_ = *K*_*D*_ = 0 *in Equation* (1), *the following integral feedback is deduced:*

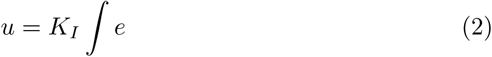

*Compare Equation* (2) *with the references above on homeostasis, and Somvanshi et al. (2015). See Abouaïssa et al. (2017b), and the references therein, for an application to ramp metering on freeways in order to avoid traffic congestion*.

Conditions for DC have already been investigated by several authors: Karin *et al*. (2017a,b); Sontag (2017); Villaverde *et al*. (2017). Parameter identification, which plays a key rôle in most of those studies, leads, according to the own words of Karin *et al*. (2017b), to some kind of “discrepancy,” which is perhaps not yet fully cleared up. We suggest therefore another roadmap, *i.e., intelligent* feedback controllers as defined by Fliess *et al*. (2013). Many concrete applications have already been developed all over the world. Some have been patented. The bibliography contains for obvious reasons only recent works in biotechnology: Bara *et al*. (2016); Join *et al*. (2017a); Lafont *et al*. (2015); MohammadRidha *et al*. (2018); Tebbani *et al*. (2016).^2^

An unexpected byproduct is derived from Remark 1.1 and the comparison in Abouaïssa *et al*. (2017b) between Equation (2) and our *intelligent proportional* controller (Fliess *et al*. (2013)). Exact adaptation means now a “satisfactory” behavior thanks to the feedback loop defined by Equation (2) when the working conditions are “mild.” The result by Karin *et al*. (2016) via a mechanism for DC based on known hormonal circuit reactions, which states that exact adaptation does not guarantee dynamical compensation, remains therefore valid in this new context.

This exploratory research report is organized as follows. Intelligent controllers are summarized in Section 2, where the connection between classic PIs and intelligent proportional controllers is also presented. Section 3, which is hevily influenced by Abouaïssa *et al*. (2017b), defines dynamic compensation and exact adaptation. Section 4 displays various convincing, but academic, computer experiments. Some concluding remarks may be found in Section 5.

## 2. Model-free control and intelligent controllers

See Fliess *et al*. (2013) for full details.

### 2.1. Generalities

#### 2.1.1. The ultra-local model

The poorly known global description of the plant, which is assumed for simplicity’s sake to be SISO (single-input single output),^3^ is replaced by the *ultra-local model*:

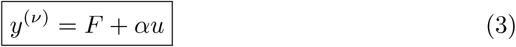

where:

- the control and output variables are respectively *u* and *y*;
- the derivation order *ν* is often equal to 1, sometimes to 2; in practice *ν* ≥ 3 has never been encountered;
- the constant *α* ∈ ℝ is chosen by the practitioner such that *αu* and *y*(*v*) are of the same magnitude; therefore does not need to be precisely estimated.

The following comments might be useful:

- Equation (3) is only valid during a short time lapse and must be continuously updated;
- *F* is estimated via the knowledge of the control and output variables *u* and *y*;
- *F* subsumes not only the system structure, which most of the time will be nonlinear, but also any external disturbance.

#### 2.1.2. Intelligent controllers

Set, *v* = 2. Close the loop with the following *intelligent proportional-integral-derivative controller*, or *iPID*,

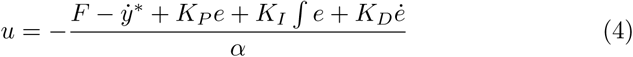

where:

- *e* = *y − y*\* is the tracking error,
- *K*_*P*_, *K*_*I*_, *K*_*D*_ ∈ ℝ are the tuning gains.

When *K*_*I*_ = 0, we obtain the *intelligent proportional-derivative controller*, or *iPD*,

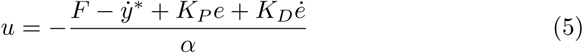

When *v* = 1 and *K*_*I*_ = *K*_*D*_ = 0, we obtain the *intelligent proportional controller*, or *iP*, which is the most important one,

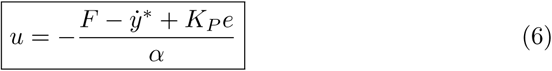

Combining Equations (3) and (6) yields:

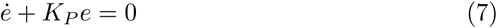

where *F* does not appear anymore. The tuning of *K*_*P*_ is therefore straightforward.

###### Remark 2.1.

*See Join et al. (2017b) for a comment on those various controllers*.

#### 2.1.3. Estimation of F

Assume that *F* in Equation (3) is “well” approximated by a piecewise constant function *F*_est_.^4^ The estimation techniques below are borrowed from Fliess *et al*. (2003, 2008) and Sira-Ramírez *et al*. (2014). Let us summarize two types of computations:

1. Rewrite Equation (3) in the operational domain (see, *e.g*., Erdélyi (1962)):

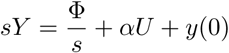

where Φ is a constant. We get rid of the initial condition *y*(0) by multiplying both sides on the left by 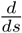

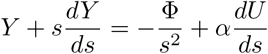

Noise attenuation is achieved by multiplying both sides on the left by *s*^−2^, since integration with respect to time is a lowpass filter. It yields in the time domain the realtime estimate, thanks to the equivalence between 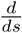 and the multiplication by *−t*,

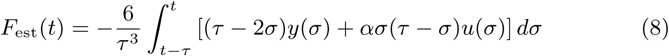

where *τ* > 0 might be quite small. This integral may of course be replaced in practice by a classic digital filter.
2. Close the loop with the iP (6). It yields:

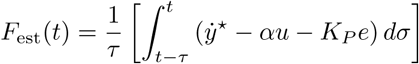

###### Remark 2.2.

*From a hardware standpoint, a real-time implementation of our intelligent controllers is also cheap and easy (Join et al. (2013))*.

### 2.2. PI and iP

Consider the classic continuous-time PI controller

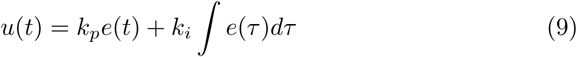

A crude sampling of the integral 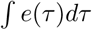 through a Riemann sum *I*(*t*) leads to

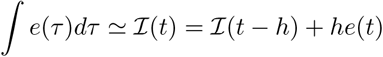

where *h* is the sampling interval. The corresponding discrete form of Equation (9) reads:

> *u*(*t*) = *k*_*p*_*e*(*t*) + *k*_*i*_*I*(*t*) = *k*_*p*_*e*(*t*) + *k*_*i*_*I*(*t − h*) + *k*_*i*_*he*(*t*)

Combining the above equation with

> *u*(*t − h*) = *k*_*p*_*e*(*t − h*) + *k*_*i*_*I*(*t − h*)

yields

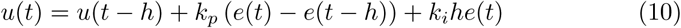

##### Remark 2.3.

*A trivial sampling of the “velocity form” of Equation* (9)

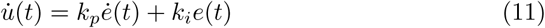

*yields*

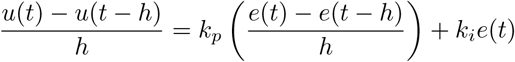

*which is equivalent to Equation* (10).

Replace in Equation (6) *F* by *ẏ*(*t*) *−αu*(*t − h*) and therefore by

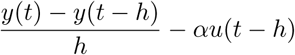

It yields

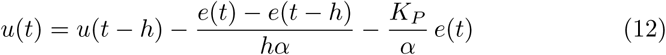

FACT.- Equations (10) and (12) become identical if we set

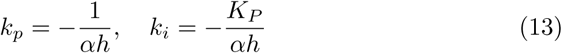

##### Remark 2.4.

*This path breaking result was first stated by d’Andréa-Novel et al. (2010):*

- *It is straightforward to extend it to the same type of equivalence between PIDs and iPDs*.
- *It explains apparently for the first time the ubiquity of PIs and PIDs in the industrial world*.

## 3. Exact adaptation and dynamic compensation

Equation (11) shows that integral and proportional-integral controllers are close when

1. *ė* remains small,
2. the reference trajectory *y*\* starts at the initial condition *y*(0) or, at least, at a point in a neighborhood,
3. the measurement noise corruption is low.

The following conditions might be helpful:

- the reference trajectory *y*\* is “slowly” varying, and starts at the initial condition *y*(0) or, at least, at a point in its neighborhood,^5^
- the disturbances and the corrupting noises are rather mild.

Then the performances of the integral controller should be decent: this is *exact adaptation*, or *homeostasis*. When the above conditions are not satisfied, *dynamic compensation* means that one at least of the feedback loops in Section 2.1 is *negative, i.e*., fluctuations around the reference trajectory due to perturbations and input changes are attenuated.^6^

## 4. Two computer experiments

The two academic examples below provide easily implementable numerical examples. They are characterized by the following features:

- *K*_*I*_ = 0.5 (resp. *K*_*I*_ = 1) for the integral feedback in the linear (resp. nonlinear) case.
- *α* = *K*_*P*_ = 1 for the the iP (6) in both cases.
- The sampling period is 10ms.
- In order to be more realistic, the output is corrupted additively by a zero-mean white Gaussian noise of standard deviation 0.01.

### 4.1. Linear case

Consider the input-output system defined by the transfer function

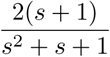

Several reference trajectories are examined:

i. Setpoint and 50% efficiency loss of the actuator: see Figures 1 see 2.
ii. Slow connection between two setpoints: see Figures 3 and 4.
iii. Fast connection: see Figures 5 and 6.
iv. Complex reference trajectory: see Figures 7 and 8.

**Figure 1:**
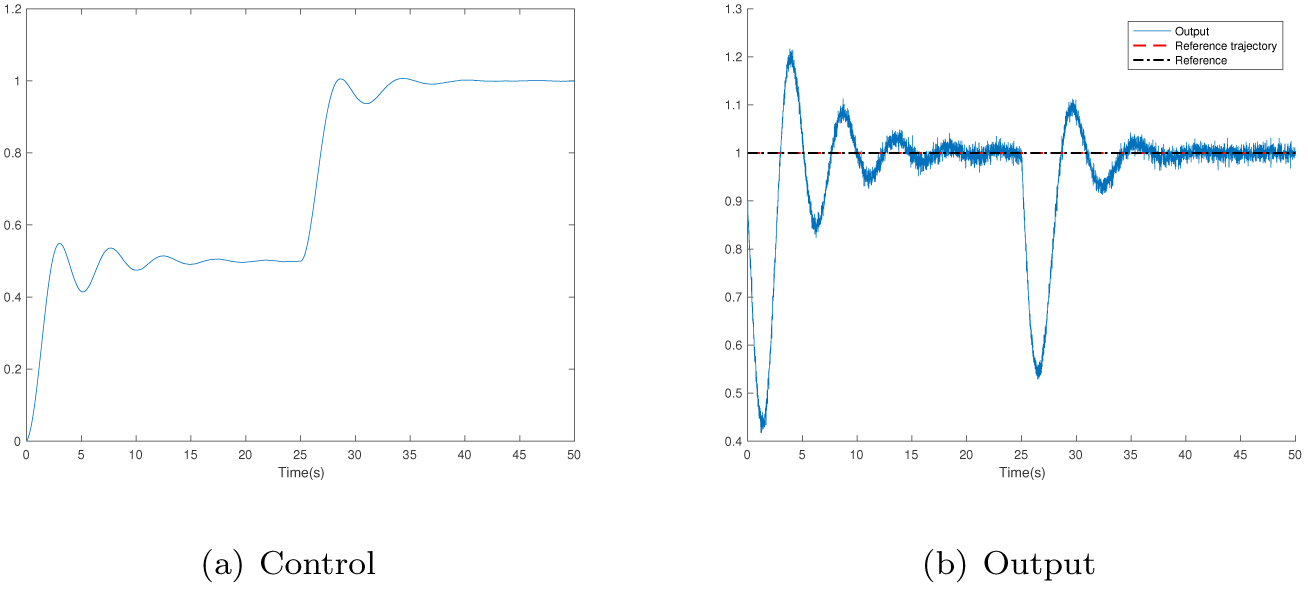
Integral feedback, constant reference trajectory, control efficiency loss

**Figure 2:**
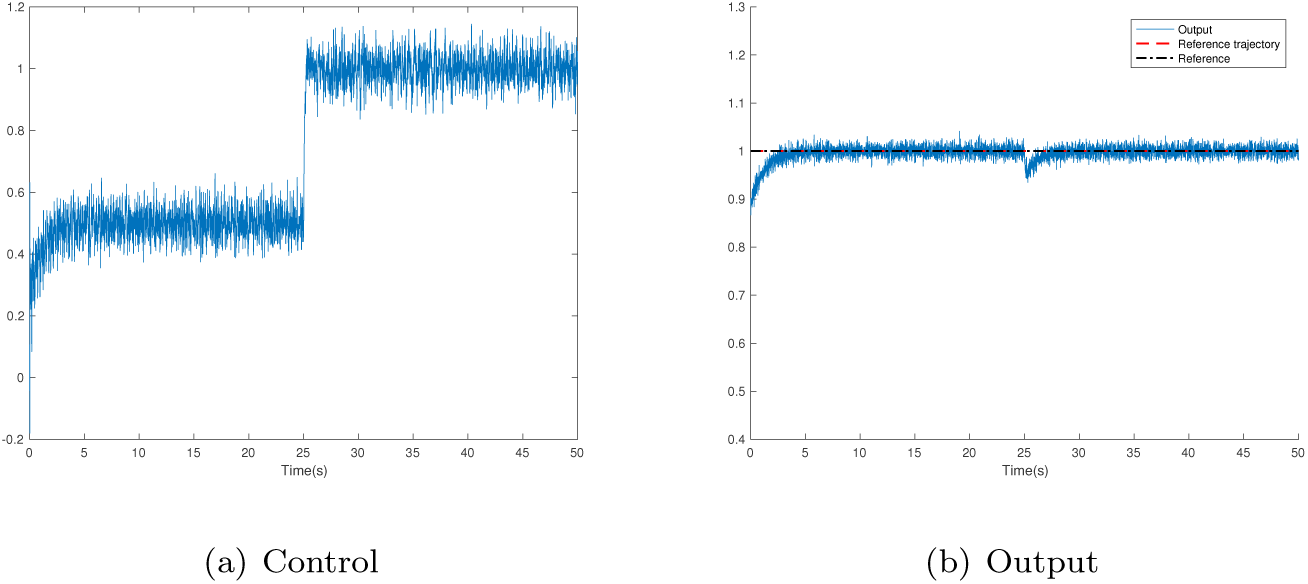
iP, constant reference trajectory, control efficiency loss

**Figure 3:**
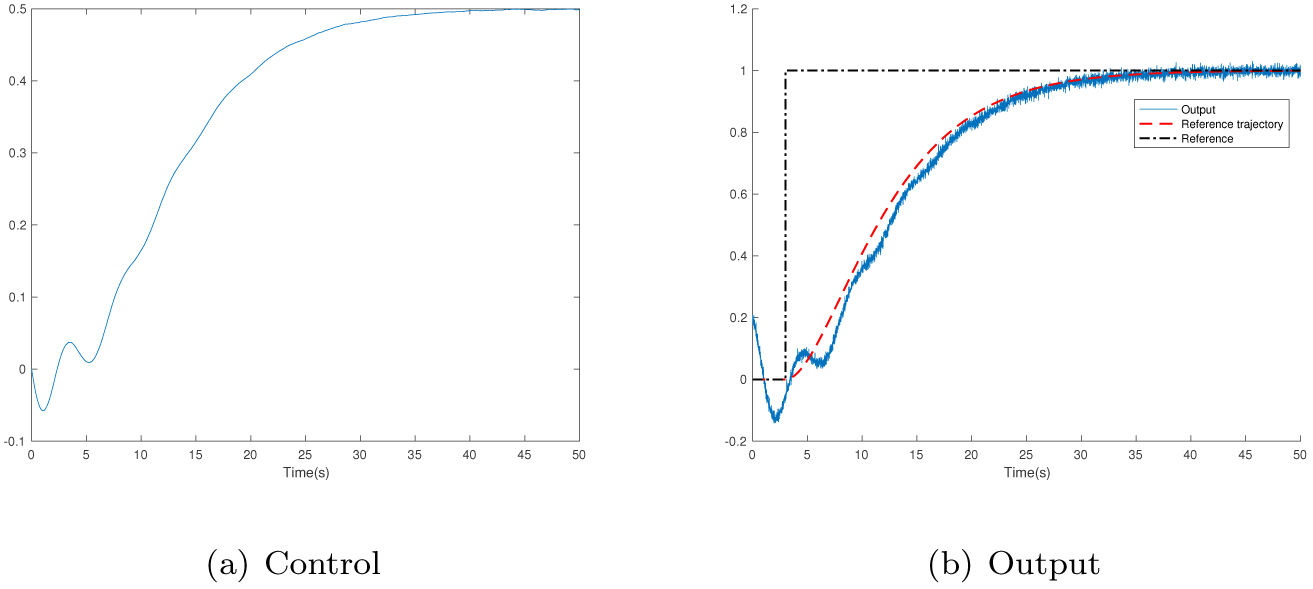
Integral feedback, slow connection

**Figure 4:**
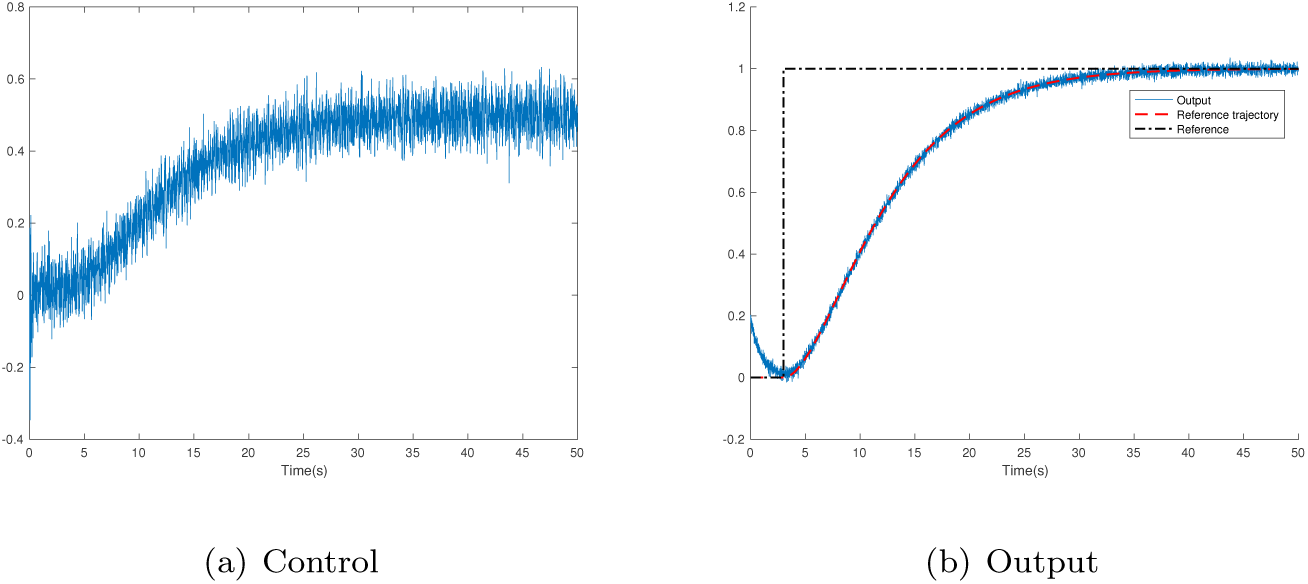
iP, slow connection

**Figure 5:**
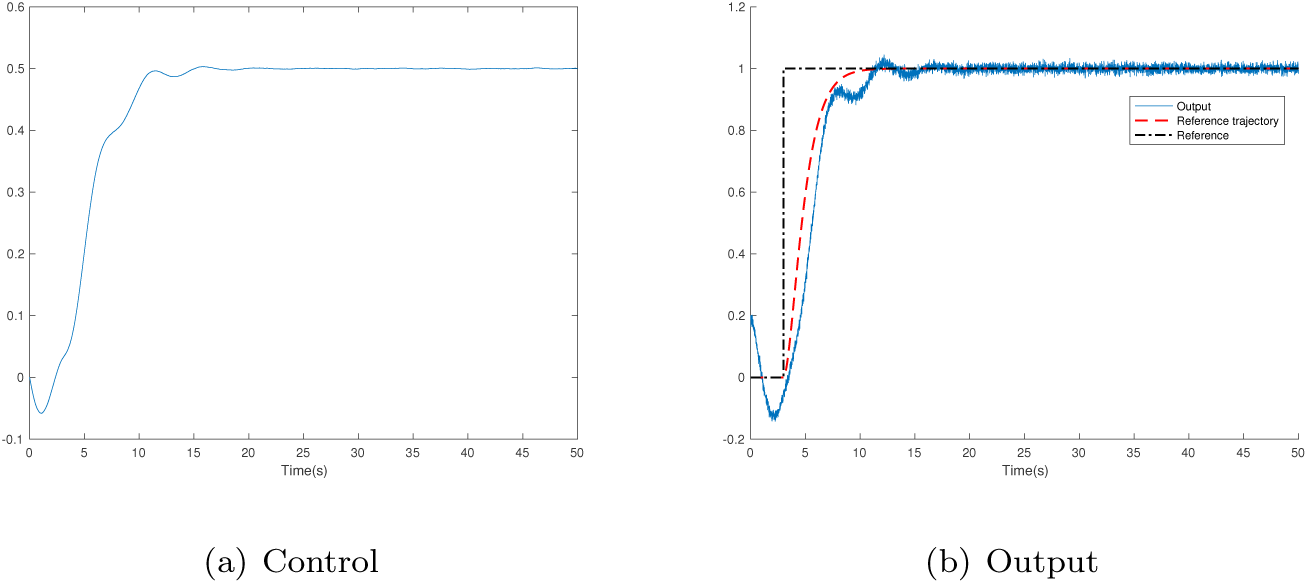
Integral feedback, fast connection

**Figure 6:**
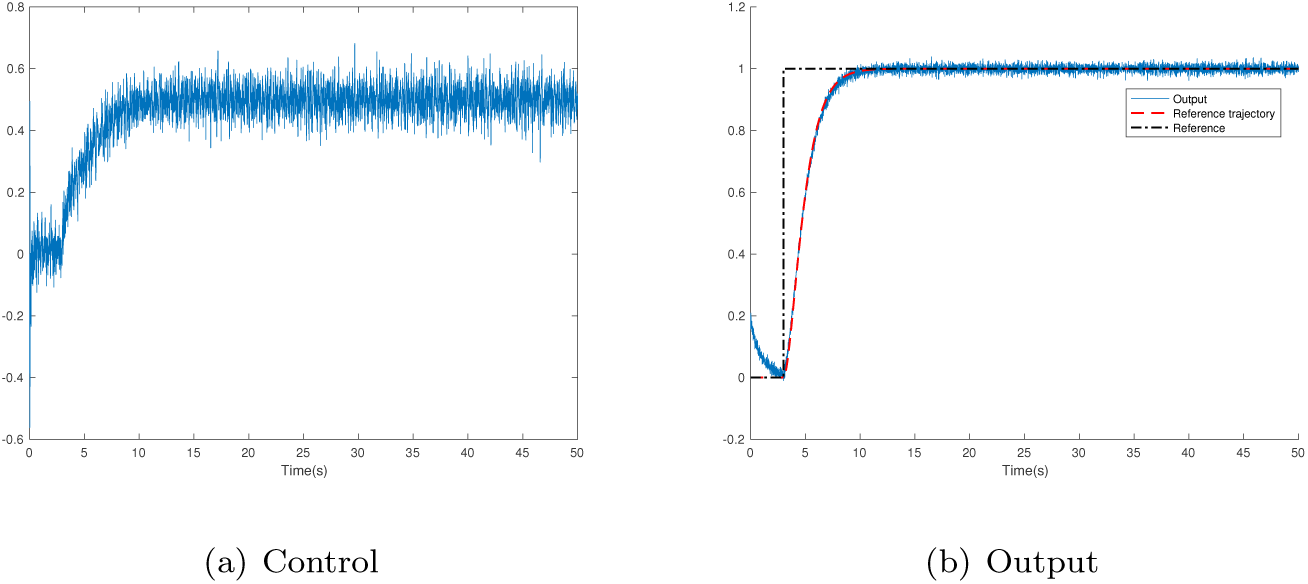
iP, fast connection

**Figure 7:**
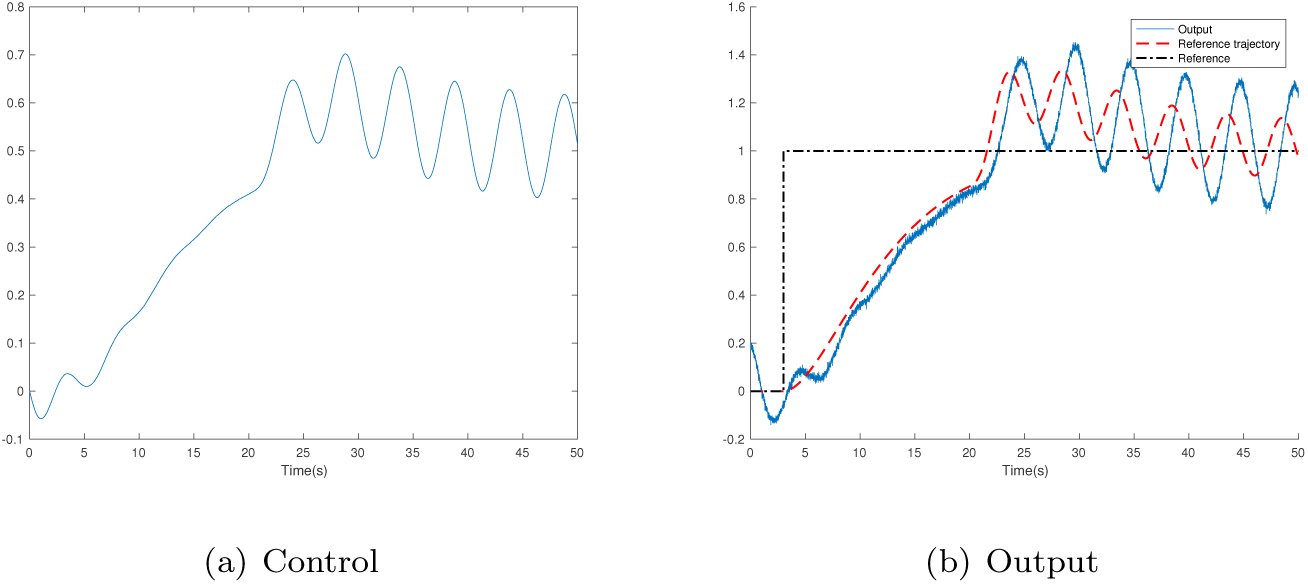
Integral connection, complex reference trajectory

**Figure 8:**
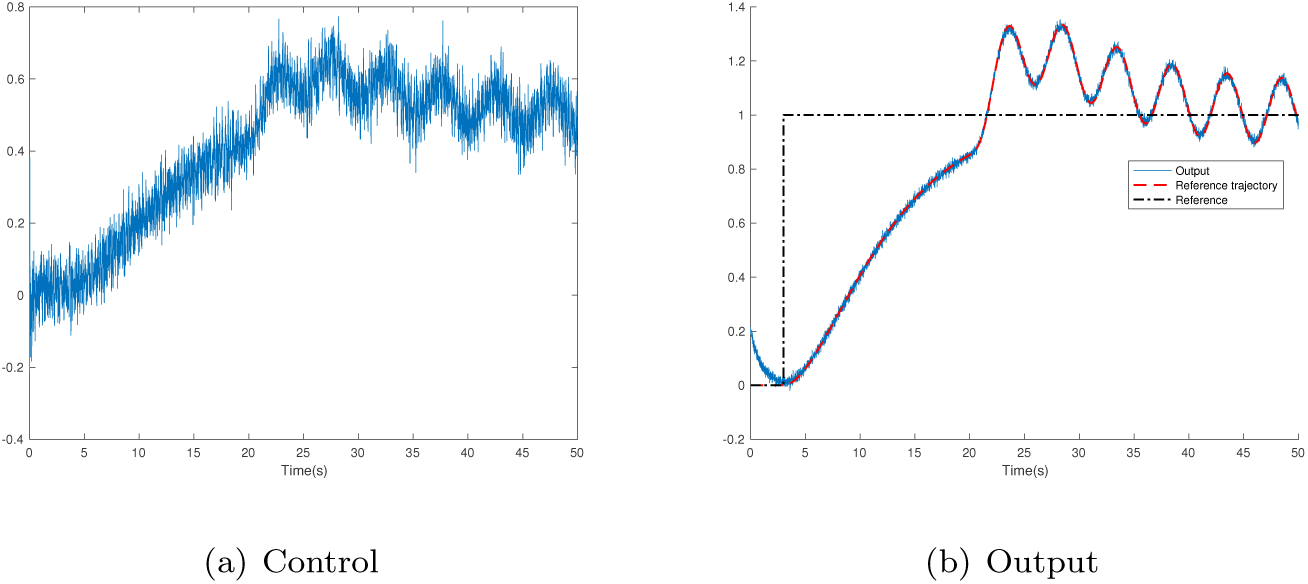
iP, complex reference trajectory

In the first scenario, the control efficiency loss is attenuated much faster by the iP than by the integral feedback. The behaviors of the integral feedback and the iP with respect to a slow connection are both good and cannot be really distinguished. The situation change drastically with a fast connection and a complex reference trajectory: the iP becomes vastly superior to the integral feedback. Exact adaptation works well only in the second scenario, whereas dynamic compensation yields always excellent results.

### 4.2. Nonlinear case

Consider

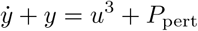

where *P*_pert_ is a perturbation. Introduce three scenarios:
i. Setpoint without any perturbation, *i.e., P*_pert_ = 0: see Figures 9 and 10.
ii. Setpoint with a sine wave perturbation which starts at *t* = 25s, *i.e*., 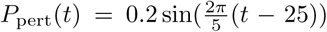 if *t* ≥ 25s, and *P*_pert_(*t*) = 0 if *t* ≤ 25s: see Figures 11 and 12.
iii. Non-constant reference trajectory without any perturbation, *i.e., P*_pert_ = 0: see Figures 13 and 14.

**Figure 9:**
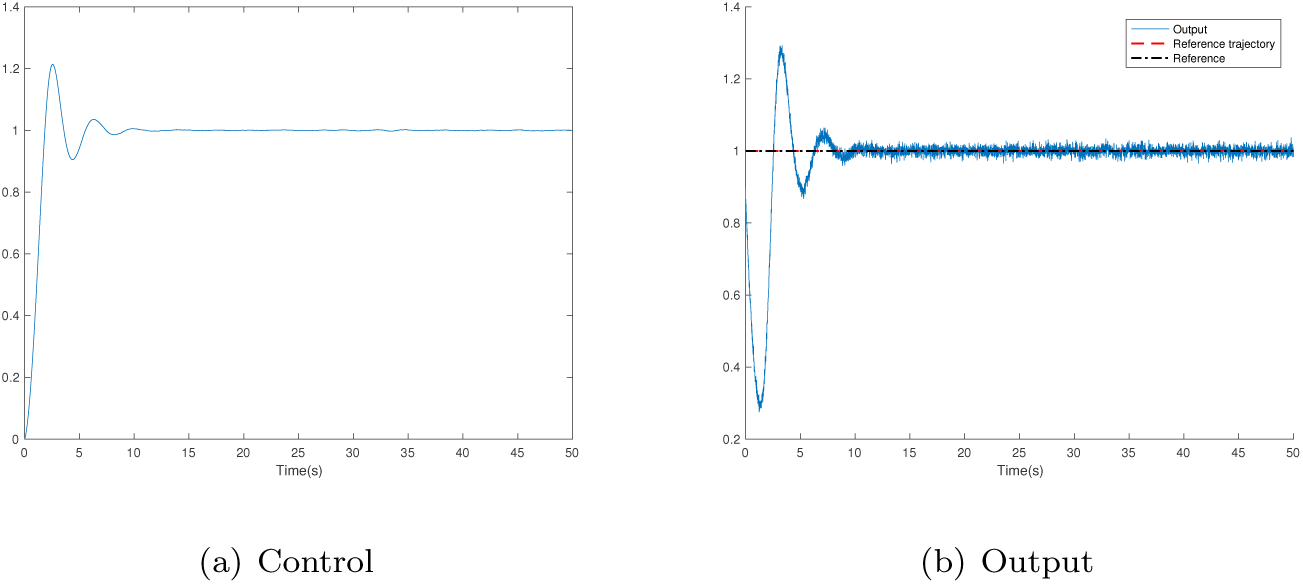
Integral feedback, constant reference trajectory, without any perturbation

**Figure 10:**
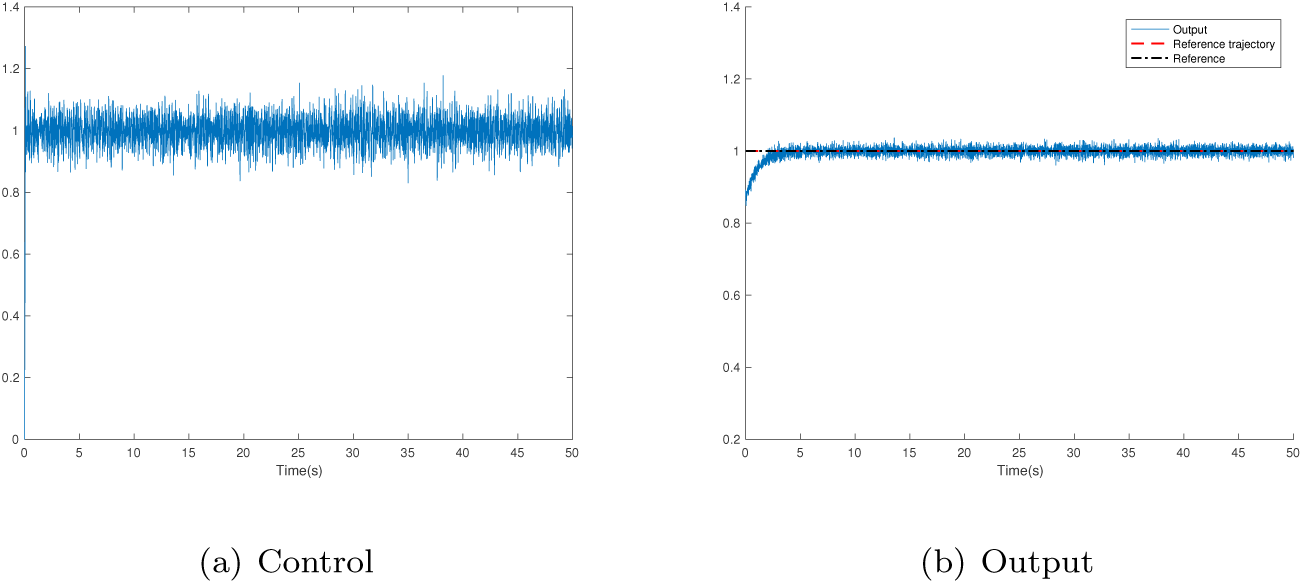
iP, constant reference trajectory, without any perturbation

**Figure 11:**
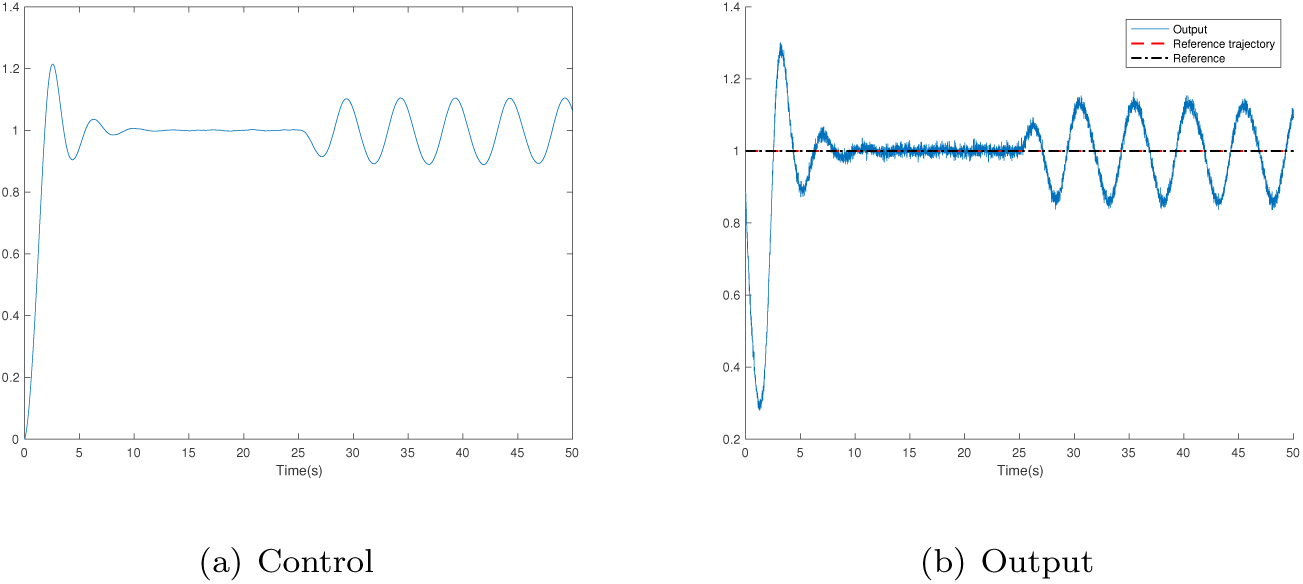
Integral feedback, constant reference trajectory, with perturbation

**Figure 12:**
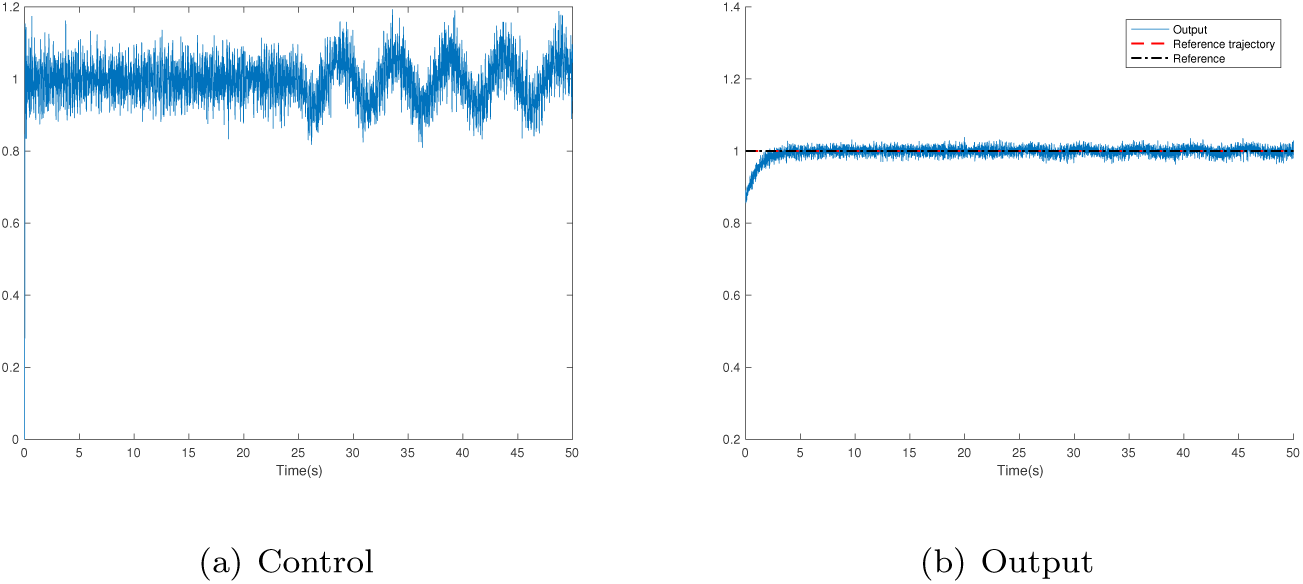
iP, constant reference trajectory, with perturbation

**Figure 13:**
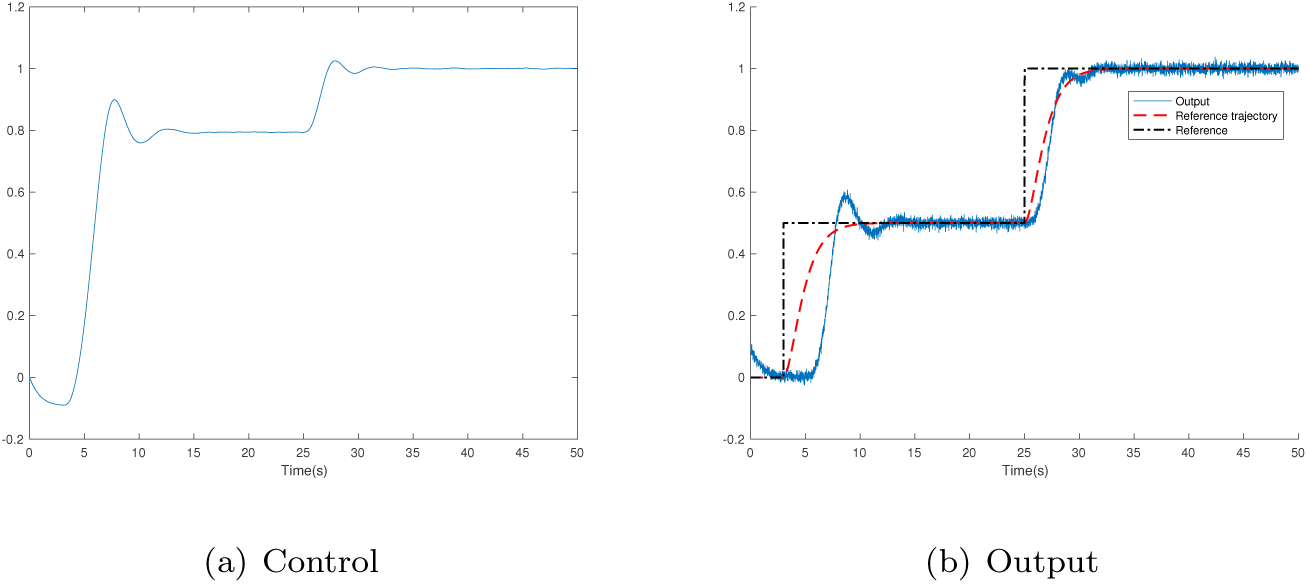
Integral feedback, non-constant reference trajectory, without any perturbation

**Figure 14:**
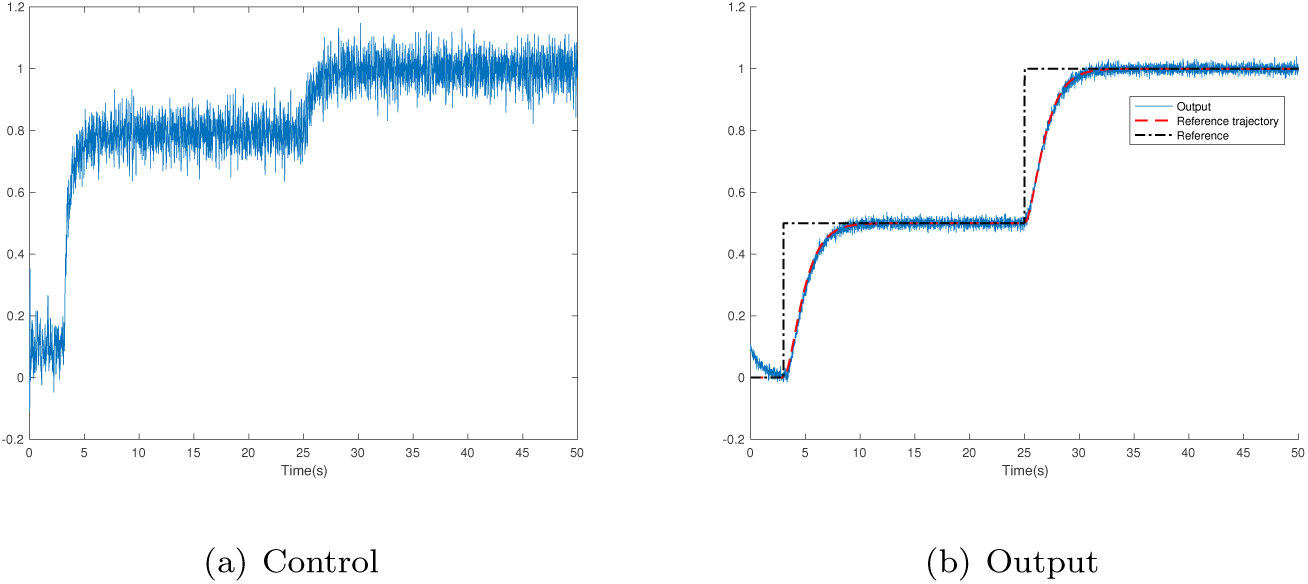
iP, non-constant reference trajectory, without any perturbation

A clear-cut superiority of the iP with respect to the integral feedback is indisputable. The behavior of dynamic compensation (resp. exact adaptation) is always (resp. never) satisfactory.

##### Remark 4.1.

*Do not believe that integral feedbacks are never adequate if non-linearities occur. See*

- *an example related to ramp metering in Abouaïssa et al. (2017b)*,
- *theoretical investigations in Sontag (2010)*.

## 5. Conclusion

In order to be fully convincing, this preliminary annoucement on homeostasis extensions needs of course to exhibit true biological examples. In our context noise corruption is also a hot topic (see, *e.g*., Briat *et al*. (2016); Sun *et al*. (2010)). The estimation and identification techniques sketched in Section 2.1 might lead to a better understanding (see also Fliess (2006, 2008)).

1 This is of course less the case in *synthetic biology, i.e*., a blending between biology and engineering (see, *e.g*., Del Vecchio *et al*. (2016) and the references therein).

2 A rather comprehensive bibliography of concrete applications is provided by Abouaïssa *et al*. (2017a).

3 For a multivariable extension, see, *e.g*., Lafont *et al*. (2015); Menhour *et al*. (2017).

4 This is a weak assumption (see, *e.g*., Bourbaki (1976)).

5 Setpoints are therefore recovered.

6 The wording “negative feedback” is today common in biology, but, to some extent, neglected in engineering, where it was quite popular long time ago (see, *e.g*., Küpfmüller *et al*. (2017)). Historical details are given by Zeron (2008) and Del Vecchio *et al*. (2015).

7 The first edition was published in 1932. Karl Küpfmüller was its sole author. He died in 1977.

## References

Abouaïssa H., Alhaj Hasan O., Join C., Fliess M., Defer D. (2017a). Energy saving for building heating via a simple and efficient model-free control design: First steps with computer simulations. 21st Int. Conf. Syst. Theor. Contr. Comput., Sinaia. https://hal.archives-ouvertes.fr/hal-01568899/en/

Abouaïssa H., Fliess M., Join C. (2017b). On ramp metering: Towards a better understanding of ALINEA via model-free control. Int. J. Contr., 90, 1018–1026.

Alon U. (2006). An Introduction to Systems Biology: Design Principles of Biological Circuits. Chapman and Hall.

Alon U., Surette M.G., Barkai N., Leibler S. (1999). Robustness in bacterial chemotaxis. Nature, 397, 168–171.

d’Andréa-Novel B., Fliess M., Join C., Mounier H., Steux B. (2010). A mathematical explanation via “intelligent” PID controllers of the strange ubiquity of PIDs. 18th Medit. Conf. Contr. Automat., Marrakech. https://hal.archives-ouvertes.fr/inria-00480293/en/

Åström K.J., Murray R.M. (2008). Feedback Systems: An Introduction for Scientists and Engineers. Princeton University Press.

Bara O., Fliess M., Join C., Day J., Djouadi S.M. (2016). Model-free immune therapy: A control approach to acute inflammation. Europ. Contr. Conf., Aalborg. https://hal.archives-ouvertes.fr/hal-01341060/en/

Bourbaki N. (1976). Fonctions d’une variable réelle. Hermann. English translation (2004): Functions of a Real Variable. Springer.

Briat C., Gupta A., Khammash M. (2016). Antithetic integral feedback ensures robust perfect adaptation in noisy biomolecular networks. Cell Syst., 2, 15–26.

Cowan N.J., Ankarali M.M., Dyhr J.P., Madhav M.S., Roth E., Sefati S., Sponberg S., Stamper S.A., Fortune E.S., Daniel T.L. (2014). Feedback control as a framework for understanding tradeoffs in biology. Integr. Compar. Biol., 54, 223–237.

Cosentino C., Bates D. (2011). An Introduction to Feedback Control in Systems Biology. CRC Press.

Del Vecchio D., Dy A.J., Qian Y. (2016). Control theory meets synthetic biology. J. Roy. Soc. Interface, 13: 20160380. http://dx.doi.org/10.1098/rsif.1916.0380

Del Vecchio D., Murray R.M. (2015). Biomolecular Feedback Systems. Princeton University Press.

Erdélyi A. (1962). Operational Calculus and Generalized Functions. Holt Rine-hart Winston.

Fliess M. (2006). Analyse non standard du bruit. C. R. Math., 342, 797–802.

Fliess M. (2008). Critique du rapport signal à bruit en communications numériques. Revue Afric. Recher. Informat. Math. Appli., 9, 419–429. https://hal.archives-ouvertes.fr/inria-00311719/en/

Fliess M., Join C. (2013). Model-free control. Int. J. Contr., 86, 2228–2252.

Fliess M., Sira-Ramírez H. (2003). An algebraic framework for linear identification. ESAIM Contr. Optimiz. Calc. Variat., 9, 151–168.

Fliess M., Sira-Ramírez H. (2008). Closed-loop parametric identification for continuous-time linear systems via new algebraic techniques. H. Garnier & L. Wang (Eds): Identification of Continuous-time Models from Sampled Data, Springer, pp. 362–391.

Join C., Bernier J., Mottelet S., Fliess M., Rechdaoui-Guérin S., Azimi S., Rocher V. (2017a). A simple and efficient feedback control strategy for wastewater denitrification. 20th World Congr. Int. Feder. Automat. Contr., Toulouse. https://hal.archives-ouvertes.fr/hal-01488199/en/

Join C., Chaxel F., Fliess M. (2013). “Intelligent” controllers on cheap and small programmable devices. 2nd Int. Conf. Contr. Fault-Tolerant Syst., Nice. https://hal.archives-ouvertes.fr/hal-00845795/en/

Join C., Delaleau E., Fliess M., Moog C.H. (2017b). Un résultat intrigant en commande sans modèle. ISTE OpenScience Automat., 1, 9 p. https://hal.archives-ouvertes.fr/hal-01628322/en/

Karin O., Alon U. (2017a). Biphasic response as a mechanism against mutant takeover in tissue homeostasis circuits. Molec. Syst. Biol., 13, 933.

Karin O., Alon U., Sontag E. (2017b). A note on dynamical compensation and its relation to parameter identifiability. bioRxiv doi: https://doi.org/10.1101/123489

Karin O., Swisa A., Glaser B., Dor Y., Alon U. (2016). Dynamical compensation in physiological circuits. Mol. Syst. Biol., 12: 886. DOI 10.15252/msb.19167216

Klipp E., Liebermeister W., Wierling C., Kowald A. (2016). Systems Biology (2nd ed.). Wiley-VCH.

Kremling A. (2012). Kompendium Systembiologie – Mathematische Model-lierung und Modellanalyse. Vieweg + Teubner. English translation (2014): Systems Biology. CRC Press.

Küpfmüller K., Mathis W., Reibiger, W. (2017). Theoretische Elektrotechnik – Eine Einführung (20. Auflage).^7^ Springer.

Lafont F., Balmat J.-F., Pessel N., Fliess M. (2015). A model-free control strategy for an experimental greenhouse with an application to fault accommodation. Comput. Electron. Agricult., 110, 139–149.

Menhour L., d’Andréa-Novel B., Fliess M., Gruyer D., Mounier H (2017). An efficient model-free setting for longitudinal and lateral vehicle control. Validation through the interconnected pro-SiVIC/RTMaps prototyping platform. IEEE Trans. Intel. Transport. Syst., 2017. DOI: 10.1109/TITS.1917.2699283

Miao H., Xia X., Perelson A.S., Wu H. (2011). On identifiability of nonlinear ODE models and applications in viral dynamics. SIAM Rev., 53, 3–39.

MohammadRidha T., Aït-Ahmed M., Chailloux L., Krempf M., Guilhem I., Poirier J.-Y., Moog C.H. (2018). Model free iPID control for glycemia regulation of type-1 diabetes. IEEE Trans. Biomed. Engin., 65, 199–206.

O’Dwyer A. (2009). Handbook of PI and PID Controller Tuning Rules (3rd ed.). Imperial College Press.

Sira-Ramírez H., García-Rodríguez C., Cortès-Romero J., Luviano-Juárez A. (2013). Algebraic Identification and Estimation Methods in Feedback Control Systems. Wiley.

Somvanshi P.R., Patel A.K., Bhartiya S., Venkatesh K.V. (2015). Implementation of integral feedback control in biological systems. Wiley Interdisc. Rev.: Syst. Biol. Med., 7, 301–316.

Sontag E.D. (2010). Remarks on feedforward circuits, adaptation, and pulse memory. IET Syst. Biol., 4, 39–51.

Sontag E.D. (2017). Dynamic compensation, parameter identifiability, and equivariances. PLoS Comput. Biol., 13: e1005447. doi: 10.1371/journal.pcbi.1005447

Sun L., Becskei A. (2010). The cost of feedback control. Nature, 467, 163–164.

Stelling J., Sauer U., Szallasi Z., Doyle F.J., Doyle J. (2004). Robustness of cellular functions. Cell, 118, 675–685.

Tebbani S., Titica M., Join C., Fliess M., Dumur D. (2016). Model-based versus model-free control designs for improving microalgae growth in a closed photobioreactor: Some preliminary comparisons. 24th Medit. Conf. Contr. Automat., Athens. https://hal.archives-ouvertes.fr/hal-01312251/en/

Villaverde A.F., Banga J.R. (2017). Dynamical compensation and structural identifiability of biological models: Analysis, implications, and reconciliation. PLoS Comput. Biol., 13, e1005878. https://doi.org/10.1371/journal.pcbi.1005878

Yi T.-M., Huang Y., Simon M.I., Doyle J. (2000). Robust perfect adaptation in bacterial chemotaxis through integral feedback control. Proc. Nat. Acad. Sci., 97, 4649–4653.

Zeron E.S. (2008). Positive and negative feedback in engineering and biology. Math. Model. Nat. Phenom., 3, 67–84.

